# Anti-aging food that improves markers of health in senior dogs by modulating gut microbiota and metabolite profiles

**DOI:** 10.1101/324327

**Authors:** Eden Ephraim Gebreselassie, Matthew I. Jackson, Maha Yerramilli, Dennis E. Jewell

**Affiliations:** Hill’s Pet Nutrition, Topeka, Kansas, United States of America; IDEXX Laboratories Inc., Westbrook, Maine, United States of America

## Abstract

Dysbiosis is one of the major changes in aging that leads to an accumulation of toxic microbial metabolites. The aim of this study was to evaluate the effect of a test food containing components of citrus, carrot, spinach and tomato on gut microbiota and age-related metabolites in senior dogs. The study was conducted on 36 dogs between 8 and 13 years of age. All dogs were maintained on a control food (control 1), which used corn as major source of fiber. After 30 days, the dogs were divided into two groups of 18 dogs. One of the groups received the test food for 30 days while the other group received the control 2 food, containing multiple whole grains as the test food but without the above added sources of fiber present in the test food. After a washout period on the control 1 food for 30 days, a cross-over was performed so that the test or the control 2 food was fed for 30 days to those dogs which had not yet been fed that food. Samples from feces and blood were collected after each 30 days period to analyze changes in gut microbial composition and metabolites. The consumption of the test food led to increased proportions of *Adlercreutzia*, *Oscillospira*, *Phascolarcobacteria*, *Faecalibacterium* and *Ruminococcus*, *Christensenellaceae*, *Ruminococcaceae*, *Cyanobacteria* and *Acidobacteria* and decreased proportions of *Megamonas*, *Salmonella*, *Enterobacteriaceae* and *Fusobacterium*. Pets had higher levels of glycerol and fatty acids and lower levels of pyrraline and mucin amino acids in feces. The test food also reduced circulating levels of pyrraline, symmetric dimethylarginine and phenolic uremic toxins, including the microbial brain toxin, 4-ethylphenyl sulfate. *Christensenellaceae* abundance was strongly associated with the observed health benefits. Fermentable fibers from fruits and vegetables enhance health in senior dogs by modulating the gut bacteria and metabolites involved in aging, kidney, brain and gut health.

## Introduction

Aging is associated with shifts in the composition of gut microbiota. An example of this is the increase in the number of facultative anaerobes and a decline in the proportion of beneficial bacteria associated with aging (1, 2). This shift in the microbial composition leads to the accumulation of toxic microbial metabolites in the body causing inflammation, oxidative stress and contributing to various diseases prominent in the aging condition (3). The reduction in the proportion of beneficial bacteria may lead to constipation, mal-absorption and longer colonic transit time. Decreased absorption of dietary protein in the upper intestine and longer colonic transit times encourage increased abundance of proteolytic bacteria, whose fermentation products deteriorate intestinal barrier integrity (4).

Foods containing fermentable fibers are known to benefit dogs by increasing nutrient absorption and reducing enteric infection (5). In an *in vitro* study, Swanson et al. (6) confirmed the fermentability of fruits and vegetables by canine fecal microflora with the resulting production of short chain fatty acids. This study evaluates the effect of a test food containing components of citrus, carrot, spinach and tomato on the microbial composition as well as metabolites associated with aging, kidney, brain and gut health in senior dogs. A recent study by Hall et al (7) showed that the consumption of a food with similar composition as the test food employed in the current study led to improvement of markers of kidney health in geriatric dogs with early stage kidney disease. This study was designed to evaluate changes in fecal microbial composition and age-related markers of health attributed to the consumption of the test food by healthy senior dogs.

## Materials and methods

### Dogs

All study protocols were reviewed and approved by the Institutional Animal Care and Use Committee, Hill’s Pet Nutrition, Inc., Topeka, KS, USA. Criteria for inclusion were healthy dogs above the age of 7 years. Dogs having chronic disease conditions such as inflammatory bowel disease, dermatitis, food allergy, cancer, tumor, kidney disease, liver disease and chronic urinary tract infections were excluded from the study. A total of 36 dogs between the ages of 8 and 13 years were grouped into a two groups of 18 each. Each group contained equal number of female and male dogs. All dogs were spayed or neutered. A summary of the description of the dogs included in this study is shown in Table 1.

**Table 1:** Description of dogs used in the study.

|  |  |
| --- | --- |
| Species | Dogs |
| Age | Group 1: $10.6 \pm 1.3$; Group 2: $10.2 \pm 1.1$ |
| Sex | Control: 9M, 9F, Test: 9M, 9F |
| Breed | Beagles |
| Initial body weight | Control: $11.2 \pm 2.1$ Kg, Test: $11.5 \pm 1.8$ Kg |
| Reproductive status | All dogs were spayed or neutered |
| Health status | Healthy |

### Foods

The study used a test food and two control foods; all in dry form. All foods were produced by Hill’s Pet Nutrition, Inc. Topeka, KS and were essentially isocaloric with respect to metabolizable energy (control 1 = 3674 kcal/kg; control 2 = 3666 kcal/kg; test = 3684 kcal/kg). The foods were formulated to meet similar nutrient profiles (Table 2) and contained grain sources such as rice, millet, oat groats, corn, wheat and/or barley. The test food contained added fiber sources from citrus, carrot, tomato and spinach in addition to the multiple grains. Unlike the test food, the first control (control 1) and the second control (control 2) foods did not have the unique fiber sources from fruit and vegetables. The first control food (control 1) used corn as major source of grain fiber and did not have multiple grain sources as the test or the control 2 food. The composition of the foods expressed as percentage of food as fed is shown in Table 2. Food analytical measurements were determined by Eurofins Scientific Inc. (Des Moines, IA) using Association of Analytical Communities (AOAC) methods.

**Table 2.** Comparison of the three different foods as fed (g/100g)

| **Nutrient** | **Control 1** | **Control 2** | **Test Food** |
| --- | --- | --- | --- |
| Moisture | 7.6 | 8.91 | 9 |
| Ash | 4.52 | 4.8 | 4.41 |
| Crude Fiber | 1.2 | 2 | 2.5 |
| Crude Protein | 19.6 | 17.77 | 19.89 |
| Carbohydrates* | 54.11 | 53.64 | 48.92 |
| Soluble Fiber | 0.5 | 1.8 | 2.7 |
| Insoluble Fiber | 6.6 | 7.2 | 5.8 |
| Crude Fat | 12.67 | 12.98 | 15.18 |
| C18:2 Omega 6 (Linoleic) | 3.43 | 3.29 | 3.32 |
| C18:3 omega 3 (alpha-Linolenic) | 0.36 | 0.41 | 0.75 |
| C20:4 Omega 6 | 0.05 | 0.05 | 0.09 |
| C20:5 EPA Omega 3 | 0.01 | 0.01 | 0.08 |
| C22:6 DHA Omega 3 | 0.01 | 0.01 | 0.06 |
| Omega 3 Sum | 0.4 | 0.43 | 0.92 |
| Omega 6 Sum | 3.54 | 3.41 | 3.49 |
| C16:1 Palmitoleic | 0.28 | 0.26 | 0.26 |
| C18:0 Stearic | 0.93 | 0.91 | 0.91 |
| Lysine | 0.92 | 0.84 | 1.27 |
| Threonine | 0.7 | 0.62 | 0.76 |
| Tryptophan | 0.24 | 0.19 | 0.29 |

\*Carbohydrate (Nitrogen-free extract) = 100% - (%Protein + %Fat + %Fiber + %Ash + %Moisture)

### Study design and sample collections

All dogs were maintained on control 1 food for 30 days and were divided into two groups. At the beginning of the test food feeding period, one of the groups received the test food while the other group received control 2 food for 30 days. Both groups were then fed the control 1 food for the next 30 days after which a cross-over was performed so that the test or the control 2 food were fed for 30 days to dogs which did not eat them during the first assignment to test foods. Water was available *ad libitum*. All dogs were meal fed from electronic feeders, where fresh food was offered daily with amounts calculated to maintain body weight. Exposure to food was allowed for up to 30 minutes to complete diet consumption. Daily food intake (g/d) was recorded for each dog. Body weights were measured weekly. Blood and fecal samples were collected at the end of each 30 days period to compare the effect of food on the abundance of various bacterial genera and various metabolites (Table 3).

**Table 3.**
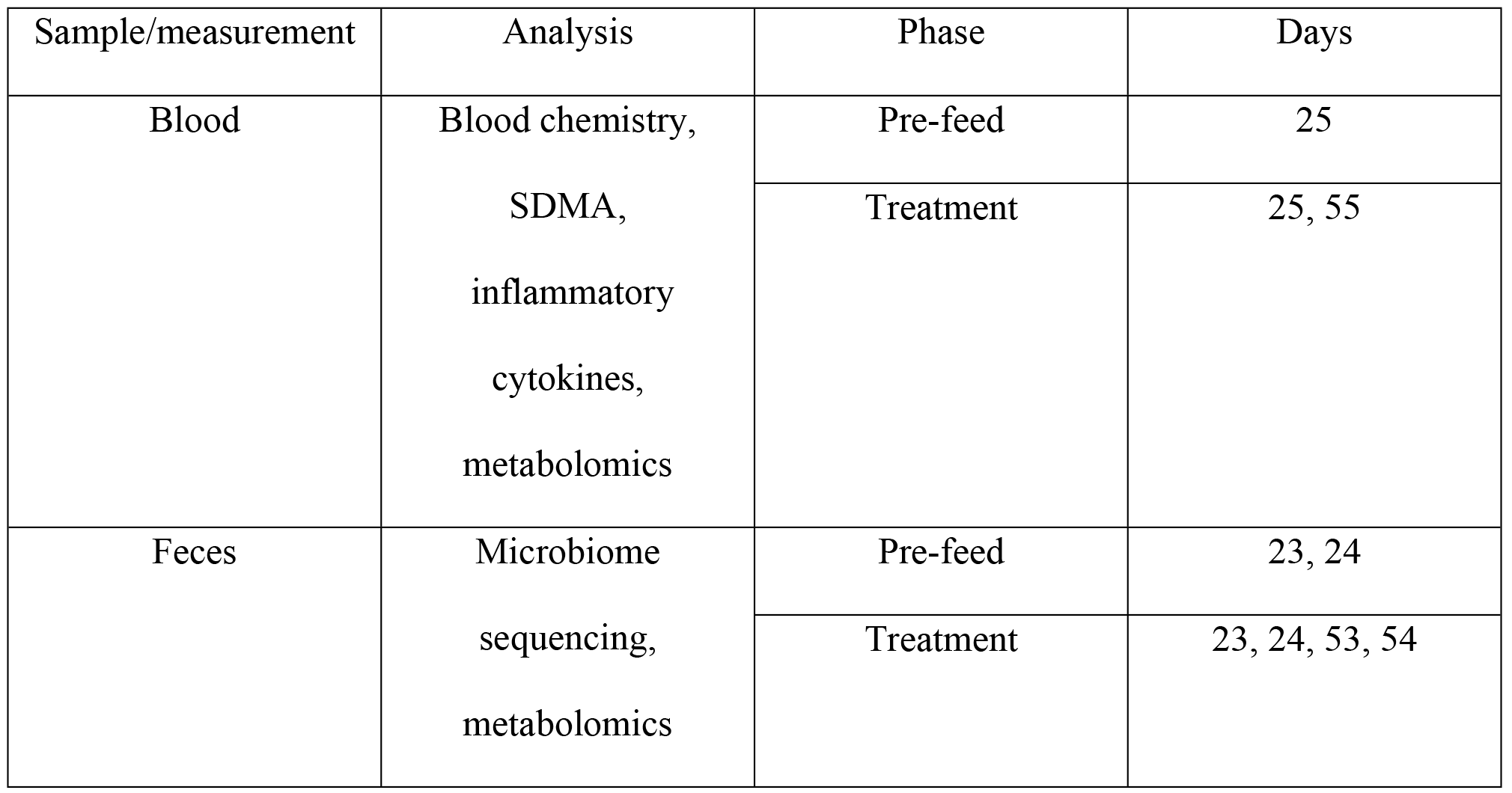
Sample analyses and measurement.

| Sample/measurement | Analysis | Phase | Days |
| --- | --- | --- | --- |
| Blood | Blood chemistry,<br>SDMA,<br>inflammatory<br>cytokines,<br>metabolomics | Pre-feed | 25 |
|  |  | Treatment | 25, 55 |
| Feces | Microbiome<br>sequencing,<br>metabolomics | Pre-feed | 23, 24 |
|  |  | Treatment | 23, 24, 53, 54 |

### Sequencing of the 16S rRNA gene

Fecal samples were collected within 30 minutes of defecation and stored at −80°C until processed. Approximately 25mg of frozen stool homogenate was used for DNA isolation with MoBio PowerFecal^®^ Kit (MoBio, Carlsbad, CA). Instructions provided by the manufacturer were followed except that a sonication step was added before vortexing the bead tubes with feces samples for 15 minutes. The DNA extracts were stored at −20°C until further processed. One microliter of each DNA sample was used to amplify the V3V4 region of the 16S rRNA gene using primers 347F and 803R containing Illumina adapters (8). Amplification was performed on BioRad C1000 Touch Thermal Cycler under the following conditions: 25 cycles of denaturation at 95°C for 30 seconds, annealing at 55°C for 30 seconds extension at 72°C for 45 seconds, and a final elongation step at 72°C for 5 minutes. An internal normalized mock community DNA and PCR-grade water were used as positive and negative controls, respectively. The mock community was formed by mixing genomic DNA of 28 bacterial species representing 25 genera obtained from the American Type Culture Collection (ATCC, Rockville, MD). The mock community represented equal copy numbers of the 16S rRNA gene of each species as described by Diaz et al. (9).

PCR amplicons (25μl) were purified by using Agencourt AmPure XP beads (Beckman Coulter) and concentrations were measured by using Qubit fluorometer 3.0 (Life Technologies). The quality of the amplicon was assessed by using Agilent 2100 Bioanalyzer. Index PCR, library quantification, normalization and pooling were performed following the Illumina’s 16S metagenomic sequencing library preparation protocol (Part # 15044223 Rev. A, Illumina, CA). Libraries were mixed with Illumina generated PhiX control library and denatured using fresh NaOH. Final sequencing libraries were then loaded onto the Illumina Miseq v3 reagent cartridge and 251-base paired-end reads were generated using Miseq Control Software (MCS) 2.4., RTA 1.18.54 and Miseq Reporter 2.4. For every Miseq run, a mock community sample and water were run as a positive and a negative control, respectively.

The reads were de-multiplexed using Miseq built-in workflow to obtain FASTQ files processed using Mothur, version 1.32 (10). Sequences were retained based on criteria such as having reads between 431 and 458 base pairs, maximum ambiguous bases of 0 and maximum homopolymer length of 6. The remaining sequences were chimera detected using the UCHIME algorithm implemented in MOTHUR and excluded from further processing (11). All retained sequences were aligned to the GreenGenes 16S rRNA gene reference database of (gg.13.5.99). The database was used for taxonomical assignment of operational taxonomic units (OTUs) at an 80% confidence threshold by using the naïve Baysian algorithm (12) implemented in MOTHUR.

### Blood and fecal metabolites

Metabolomic profiles of blood and fecal samples were determined by Metabolon (Durham, NC). The methods utilized a Waters ACQUITY ultra-performance liquid chromatography (UPLC) and a Thermo Scientific Q-Exactive high resolution/accurate mass spectrometer interfaced with a heated electrospray ionization (HESI-II) source and Orbitrap mass analyzer operated at 35,000 mass resolution. Different aliquots of sample extracts were analyzed under different chromatographic conditions optimized for hydrophilic or hydrophobic compounds (13). Standards present in each aliquot were used to ensure injection and chromatography consistency. Peaks were identified and processed using proprietary hardware and software. The relative quantification of the metabolites was performed by using area-under-the curve.

Symmetric dimethylarginine (SDMA) concentrations in blood samples were determined using liquid chromatography-mass spectroscopy (LC-MS) as described by Hall et al. (14).

### Statistical analysis

Matched-pair analyses were performed with JMP version 12 (SAS Institute, Carry, NC) to compare differences between means of the microbial abundances and relative levels of metabolites on samples collected from the same dog after the consumption of the test or the control 2 food. P values were calculated for differences between means and false discovery rate (FDR) corrections were made on each group of markers. FDR-P values less than 0.05 were considered significant. A bivariate regression analysis was performed to evaluate correlations between the changes in the microbial abundance and fecal metabolites.

## Results

### Food intake and body weight

All dogs completed the study successfully and there was no adverse health report. There was a trend (P=0.06) inthe body weights of the dogs consuming the test food (10.96 Kg) to increase when compared to the control 2 food (10.74 Kg). There was very little difference in intakes of the test (113.41 Kcal/Body weight^0.75, SE=3.89) or the control 2 food (113.9 Kcal/Body weight^0.75, SE=4.15).

### Changes in the gut microbial composition

The test food led to significant changes in the proportions of bacteria at various taxa levels. Fig 1 summarizes the log (base 2) fold changes of the taxa after the consumption of the test food compared to the control 2 food. At the phylum level, *Acidobacteria* and *Cyanobacteria* increased by 0.91 and 0.72 log fold changes, respectively. This was accompanied by a −0.65 log fold reduction in the phylum *Fusobacteria*. This was equivalent to 87% and 65% increase in *Acidobacteria* and *Cyanobacteria*, respectively, and a 66% reduction in *Fusobacteria*. At the family level, *Christensenellaceae* and *Ruminococcaceae* increased by 1.76 and 0.68 log fold changes, respectively. These were equivalent to a 138% and 60% increase in the proportions of *Christensenellaceae* and *Ruminococcaceae*, respectively, compared to their levels on the Control 2 food. On the contrary, the test food led to 1.65 log fold reduction (68%) in *Enterobacteriaceae*. At the genus level, *Adlercreutzia* and *Phascolarctobacterium* increased by 77% and 68%, respectively.

**Fig 1.**
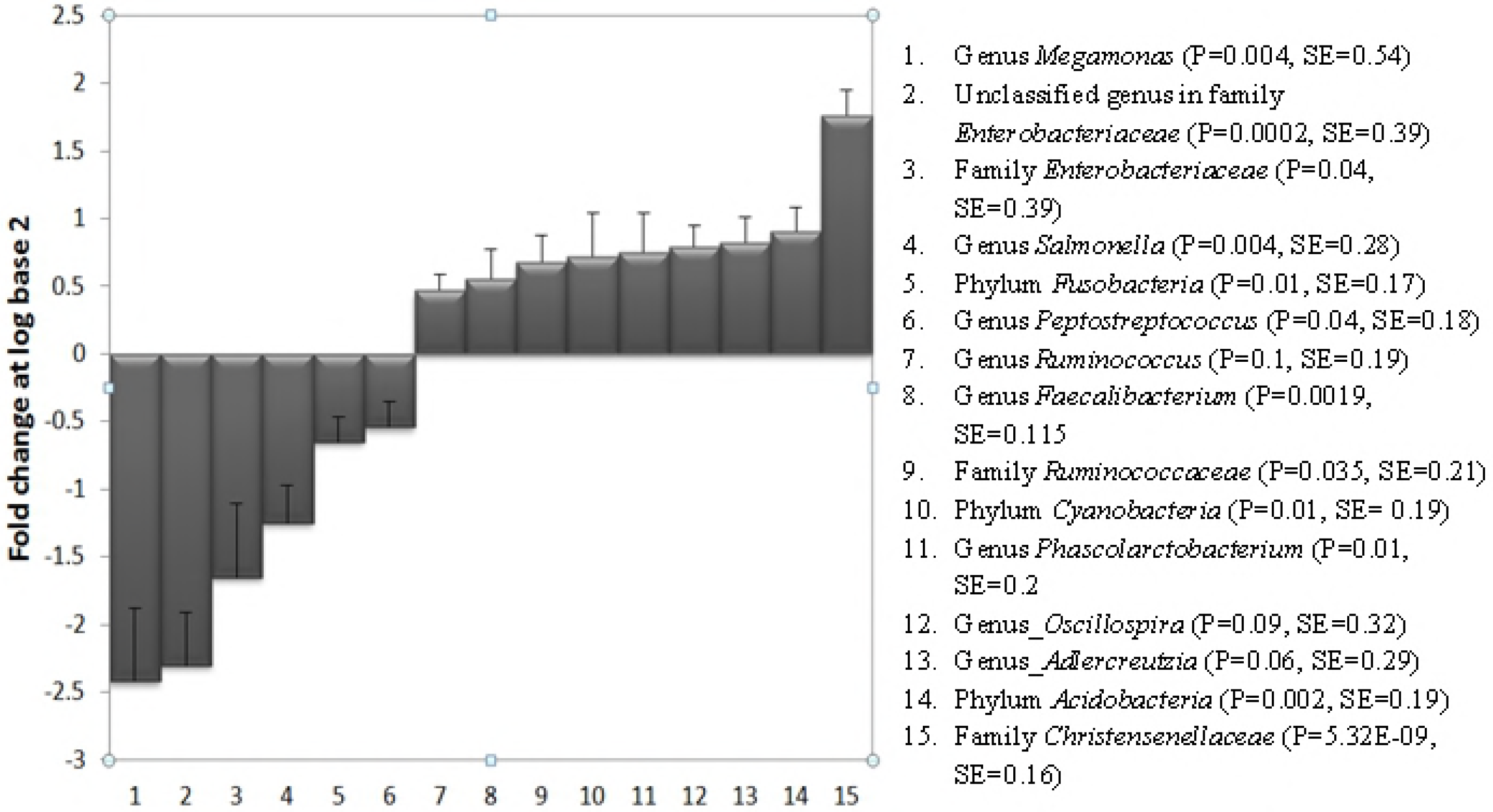
Log fold-changes (base 2) of the different taxa after the consumption of the test food. The test food led to a significant reduction in the proportion of bacteria belonging to the genus *Megamonas*, an unclassified genus in family *Enterobacteriacea*, *Salmonella* and *Peptostreptococcus*. The consumption of the test food significantly increased the proportions of the genera *Adlercreutzia*, *Oscillospira*, *Phascolarcobacterium*, *Faecalibacterium* and *Ruminococcus*. At the family level, the test food led to a significant increase in *Christensenellaceae* and *Ruminococcaceae* and a significant reduction in Enterobacteriaceae. At the phylum level, the test food increased the phyla *Acidobacteria* and *Cyanobacteria* and led to a significant reduction in *Fusobacteria*.

Although they did not meet the statistical significance, *Oscillospira* and *Ruminococcus* increased by 73% and 39%, respectively, after the consumption of the test food. This was accompanied by reductions in the proportions of the genera *Salmonella*, *Megamonas*, *Peptostreptococcus* and an unknown genus (OTU_10001 in the family *Enterobacteriaceae*) by 58%, 81%, 32%, 80%, respectively. S1 Table provides the means and standard errors of the proportions of the above taxa on the test and the control 2 foods.

### Levels of fatty acids and glycerol

The levels of fecal and circulating unsaturated fatty acids increased after the consumption of the test food (Table 4). In plasma, levels of docosahexaenoate (DHA; 24:6n3), docosapentaenoate (DPA; 22:5n6), eicosapentaenoate (EPA; 22:5n3), linolenate (18:3n3) and stearidonate (18:4n3) were increased. In feces, in addition to these fatty acids, docosapentaenoate (DPA; 22:5n3) and palmitoleate (16:1n7) were increased on the test food. Despite the similar levels of circulating glycerol on both foods, fecal levels of glycerol increased by 24% when the pets consumed the test food compared to the control 2 food (P=0.001) (Table 4).

**Table 4.** Change in levels of fatty acids and glycerol in feces and plasma of the senior dogs during the consumption of the Test versus the Control 2 food.

|  | Feces |  |  |  | Plasma |  |  |  |
| --- | --- | --- | --- | --- | --- | --- | --- | --- |
| **Metabolite in feces** | **% change on Test** | **P Value (FDR)** | **Mean difference (test-Control 2)** | **SE** | **% change on Test** | **P Value (FDR)** | **Mean difference (test-Control 2)** | **SE** |
| arachidonate (20:4n6) | -16.24 | 1.49 | -0.2 | 0.12 | 10.51 | 2.46 | 0.11 | .08 |
| docosahexaenoate (DHA; 22:6n3) | 256.47 | 3.52E-11 | 2.17 | 0.21 | 177.33 | 1.02E-10 | 1.44 | 0.14 |
| docosapentaenoate (DPA; 22:5n3) | 43.85 | 0.02 | 0.45 | 0.13 | 35.53 | 0.11 | 0.36 | 0.12 |
| docosapentaenoate (n6 DPA; 22:5n6) | 120.98 | 1.84E-07 | 1.1 | 0.15 | 57.13 | 0.0003 | 0.57 | 0.11 |
| eicosapentaenoate (EPA; 20:5n3) | 434.21 | 1.22E-13 | 3.97 | 0.3 | 191.50 | 4.83E-08 | 1.61 | 0.21 |
| glycerol | 31.70 | 0.01 | 0.36 | 0.1 | 3.33 | 1.0* | 0.04 | 0.09 |
| laurate (12:0) | -4.52 | 1.0* | -0.05 | 0.06 | 2.45 | 1.0* | 0.03 | 0.08 |
| linoleate (18:2n6) | -4.84 | 1.0* | -0.05 | 0.07 | 7.86 | 1.0* | .081 | 0.08 |
| linolenate (18:3n3 or 3n6) | 45.47 | 0.004 | 0.44 | 0.1 | 40.36 | 0.02 | 0.42 | 0.12 |
| myristate (14:0) | 10.73 | 1.0* | 0.11 | 0.07 | 5.10 | 1.0* | 0.06 | 0.1 |
| oleate/vaccenate (18:1) | 8.80 | 1.0* | 0.1 | 0.08 | 9.49 | 1.0* | .096 | 0.07 |
| palmitate (16:0) | 1.01 | 1.0* | 0.01 | 0.09 | 7.64 | 1.0* | 0.08 | 0.06 |
| palmitoleate (16:1n7) | 121.28 | 2.01E-09 | 1.23 | 0.1 | 26.21 | 0.51 | 0.28 | 0.13 |
| stearate (18:0) | -22.59 | 0.08 | -0.26 | 0.09 | 3.03 | 1.0* | 0.03 | 0.06 |
| stearidonate (18:4n3) | 776.56 | 1.33E-14 | 8.1 | 0.6 | 132.47 | 7.68E-06 | 1.29 | 0.21 |

Changes that were statistically significant ( $P < 0.05$ ) are marked grey.
1.0\*: FDR P values greater than 1 are referred as 1.

### Fecal levels of mucin amino acids

The relative fecal levels of amino acids that make up the mucin layer, such as aspartate, proline, serine and threonine were significantly affected by the type of food consumed by the senior dogs. Compared to the control 2 food, the consumption of the test food led to a 28 – 61% reduction in levels of these amino acids in the feces (Table 5).

**Table 5.** Relative levels of mucin amino acids in feces.

| Amino acid in feces | % change on Test | P Value (FDR) |
| --- | --- | --- |
| Asparagine | -61.03 | 0.007 |
| Proline | -28.5 | 0.007 |
| Serine | -36.2 | 0.004 |
| Threonine | -35.6 | 0.001 |

### Advanced glycation end products (AGE)

High levels of circulating advanced glycation end products (AGE) are associated with aging and various age-related diseases. The test food led to about 70% reductions in both circulating (P=1.15E-07 and fecal (P=5.29E-13) levels of one of the AGE, pyrraline. The circulating level of another AGE, N6-carboxymethyllysine (CML), was not affected by the different diets; but the fecal levels were higher on the test food. The third AGE, N6-carboxyethyllysine (CEL), was detected only in feces and did not change during the consumption of the different diets (Table 6).

**Table 6.** Changes in levels of advanced glycation end products (AGE)

| Matrix | AGE | % change<br>on Test | P Value<br>(FDR) | Mean difference<br>(Test-Control 2) | SE |
| --- | --- | --- | --- | --- | --- |
| Feces | Pyrraline | -69.65 | 5.29E-13 | -1.74 | 0.15 |
| Plasma | Pyrraline | -68.82 | 1.15E-07 | -1.56 | .23 |
| Feces | N6-carboxymethyllysine | 48.86 | 0.0006 | 0.44 | 0.11 |
| Plasma | N6-carboxymethyllysine | 4.25 | 0.89 | 0.05 | .07 |
| Feces | N6-carboxyethyllysine | 9.15 | 1.0 | 0.08 | .09 |

Changes that were statistically significant are marked grey.

### Changes in circulating uremic toxins

Uremic toxins are among the major toxic metabolites that lead to renal and associated diseases in aging. Some uremic toxins originate from protein fermentation in the colon by proteolytic bacteria. Products of the putrefaction process are absorbed and converted to toxic derivatives causing an increased burden on kidney function. We detected a total of 14 phenolic and indolic uremic toxins in plasma (Fig 2). The phenolic uremic toxins, 3-methyl catechol sulfate (P=0.0015, SE=0.19), 4-ethylphenyl sulfate (P=2.38E-09, SE=0.05), 3-methoxycatechol sulfate (P=0.02, SE=0.11) and 4-vinylphenol sulfate (P=0.05, SE=0.05) declined by 175%, 73%, 67% and 23%, respectively after the consumption of the test food. On the contrary, the indolic uremic toxins 5-hydroxyindole sulfate (P=2.23E-06, SE=0.05) and 7-hydroxyindole sulfate (P=2.97E-10, SE=0.06) increased by 29% and 43%, respectively, after the consumption of the test food. None of the other uremic toxins were significantly influenced by the different foods.

**Fig 2.**
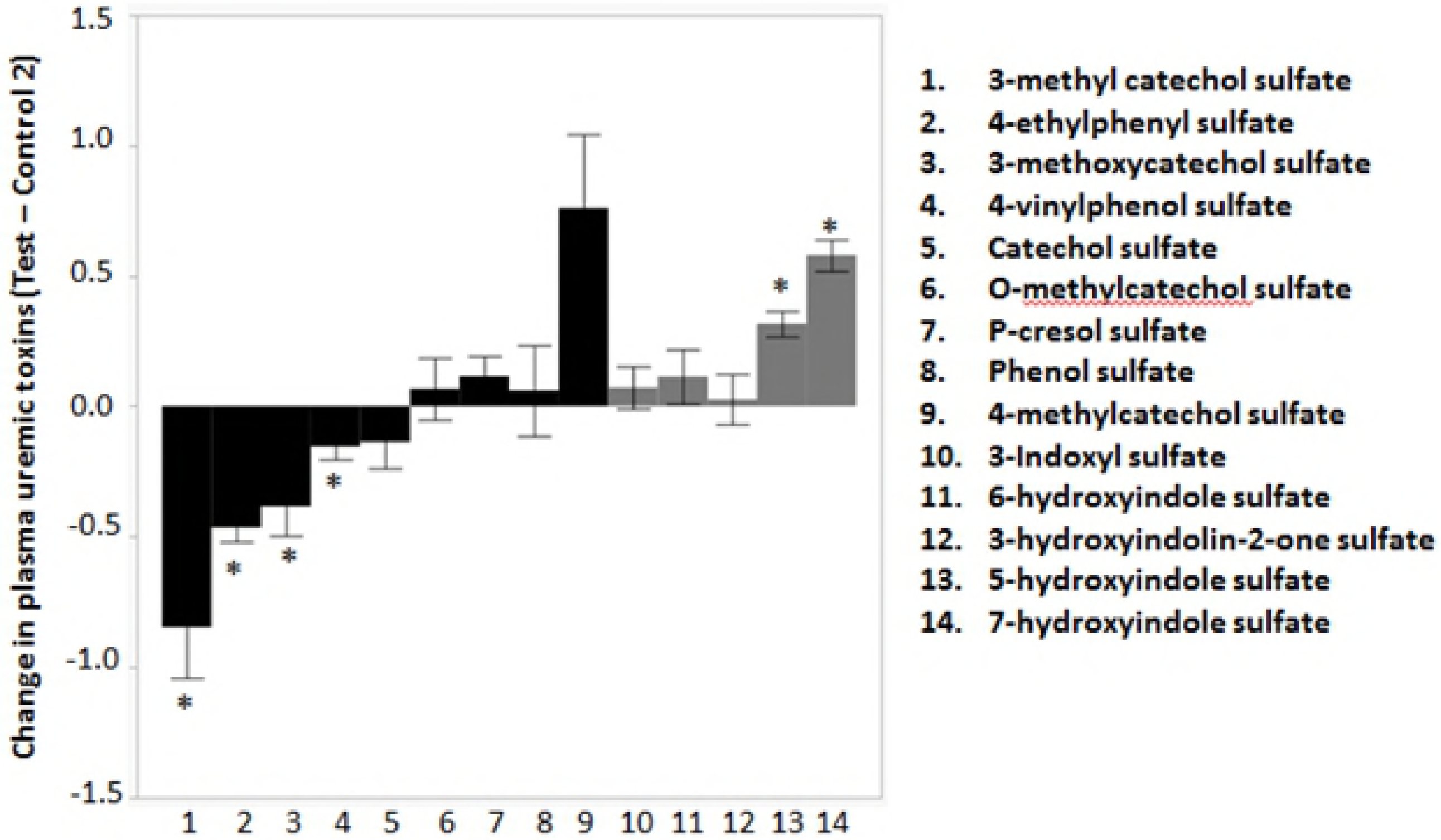
Changes in circulating levels of 9 phenolic (black bars) and 5 indolic (grey bars) uremic toxins after the consumption of the test food. The test food led to significant (*: False discovery rate corrected (FDR) P-value <0.05) reductions in levels of phenolic uremic toxins such as 3-methyl catechol sulfate, 4-ethylphenyl sulfate, 3-methoxycatechol sulfate and 4-vinylphenol sulfate. Two of the indolic uremic toxins, namely 5-hydroxyindole sulfate and 7-hydroxyindole sulfate, increased after the consumption of the test food. There were no significant changes in the typical uremic toxins such as 3-indoxyl sulfate or P-cresol sulfate.

Symmetric dimethylarginine (SDMA) is a uremic toxin originating from the host metabolism and methylation of arginine (15, 16). The test food resulted in a significant reduction in blood concentrations of SDMA (P=0.035, SE=0.2) in the senior dogs compared to the control 2 food (Fig 3).

**Fig 3.**
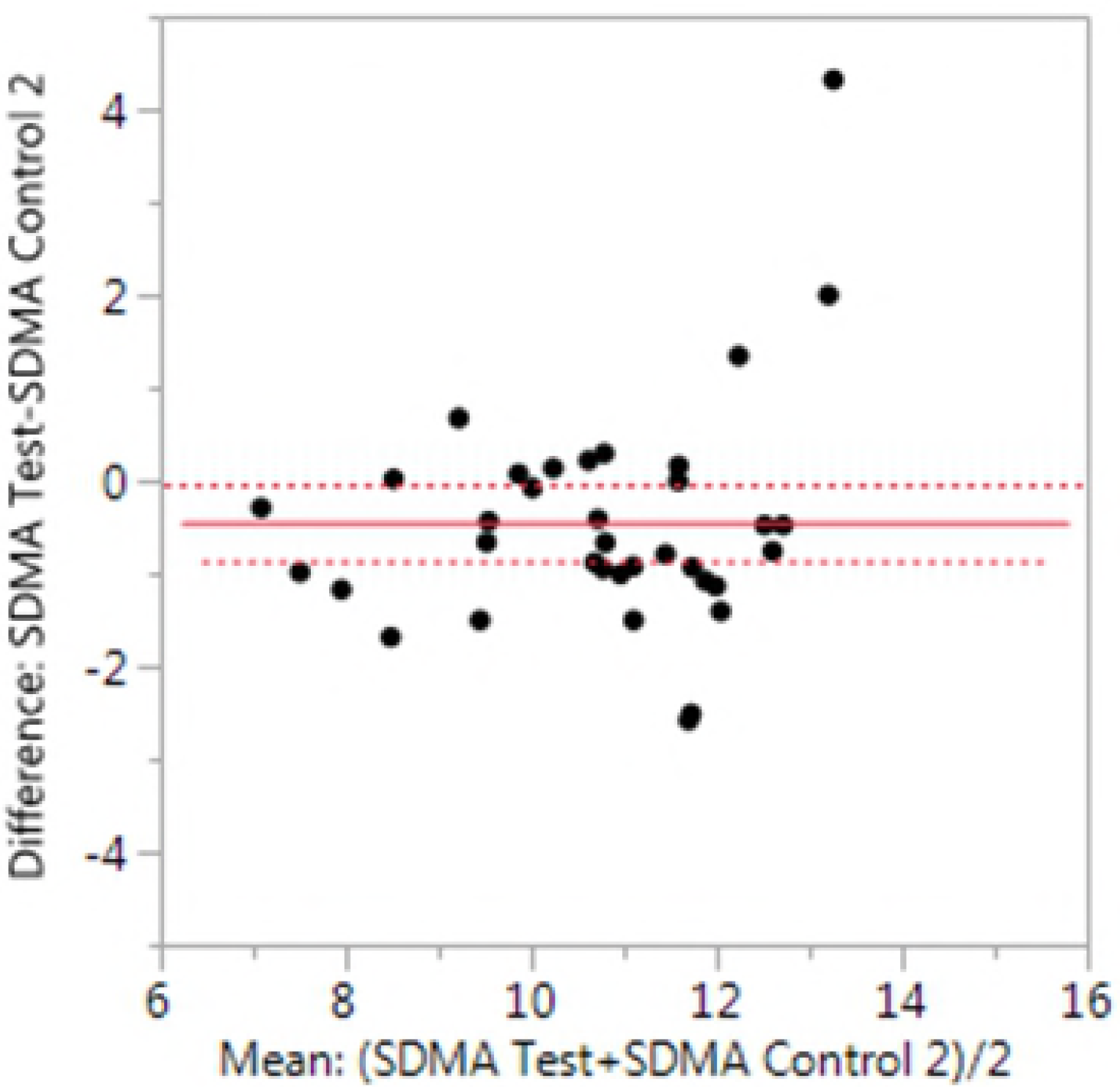
Changes in circulating levels of the renal health marker symmetric dimethylarginine (SDMA). Matched pair analyses of each dog on the test food versus the control 2 food showed significant reduction in plasma concentrations of symmetric dimethylarginine (SDMA) (P=0.035, SE=0.2) after the consumption of the test food.

### Correlations of microbial taxa with changes in metabolites

The genus *Adlercreutzi* and the family *Christensenellaceae* were strong positive predictors of glycerol levels in feces (**Table 7**). *Faecalibacterium prausnitzii*, family *Ruminococcaceae*, genus *Phascolarctobacterium* and phylum *Actinobacteria* also correlated positively with fecal levels of glycerol. On the contrary, phylum *Fusobacterium* and genus *Salmonella* negatively correlated with glycerol levels in feces (Table 7). The genera *Oscillospira* and *Adlercreutzia* also had a negative correlation with levels of pyrraline in feces. The phylum *Fusobacterium* correlated positively with fecal levels of pyrraline and threonine. *Salmonella* also had a positive correlation with pyrraline levels in feces (Table 7).

**Table 7.** Correlations of different taxa with fecal levels of glycerol, pyrraline and mucin amino acids.

| **Correlations with glycerol in feces** | **FDR P-value** | **R** |
| --- | --- | --- |
| Genus *Adlercreutzii* | 3.95E-09 | 0.51 |
| Family *Christensenellaceae* | 0.0001 | 0.42 |
| Genus *Faecalibacterium prausnitzii* | 0.02 | 0.39 |
| Family *Ruminococcaceae* | 0.02 | 0.39 |
| Genus *Phascolarctobacterium* | 0.003 | 0.33 |
| Phylum *Actinobacteria* | 0.0005 | 0.32 |
| Phylum *Fusobacterium* | 0.002 | -0.4 |
| Genus *Salmonella* | 0.005 | -0.25 |
| **Correlations with pyrrolidine in feces** |  |  |
| Family *Christensenellaceae* | 1.05E-12 | -0.52 |
| Genus *Oscillospira* | 0.002 | -0.48 |
| Genus *Adlercreutzia* | 1.52E-14 | -0.44 |
| Genus *Salmonella* | 0.026 | 0.48 |
| Phylum *Fusobacterium* | 0.017 | 0.41 |
| **Correlations with mucin amino acids in feces** |  |  |
| Family *Christensenellaceae* and threonine | 0.0004 | -0.32 |
| Family *Christensenellaceae* and serine | 0.0007 | -0.27 |
| Phylum *Fusobacterium* and threonine | 0.00015 | 0.33 |
| Phylum *Fusobacterium* and serine | 0.02 | 0.25 |

## Discussion

The test food increased the relative abundance of health-promoting bacteria belonging to the genera *Adlercreutzia* and *Phascolarctobacterium*. The abundance of the short-chain fatty acids producers *Phascolarctobacterium* was reported to have a positive association with positive mood and their number declines in elderly humans (17). Both *Adlercreutzia* and *Phascolarctobacterium* have the capacity to metabolize isoflavones to equol, which has been implicated to have antioxidative properties and prevent various age-related diseases including diabetes and obesity (18–20). Equol is also associated with a decreased risk of certain types of cancer; therefore increasing the abundance of equol-producing gut microbiota has been recommended to reduce this risk (21). In this study, although it did not reach statistical significance (P=0.17), the fecal level of equol increased by 59.6% after the consumption of the test food. Polyphenols bound to fruits and vegetables present in the test diet may have led to the increased abundance of these bacteria in the senior dogs.

The consumption of the test food led to an increase in the proportion of the butyrate producer *Faecalibacterium prausnitizii. F. prausnitzii* has been reported to have anti-inflammatory effects (22) and the abundance of *Faecalibacterium* species declines during active inflammatory bowel disease (23). The family *Christensenellacea* also increased after the consumption of the test food. Our correlation analysis showed *Christensenellacea* abundance was a strong positive predictor of fecal levels of glycerol. In a study that compared the microbiome of 416 twin-pairs, *Christensenellacea* were in a greater abundance in lean individuals compared to obese (24). *Christensenellacea* have also been reported to have the capability to produce short-chain fatty acids (SCF) (25). SCF are known to improve the intestinal barrier integrity, which is in line with the result of our correlation analysis showing a negative association between fecal levels of mucin amino acids and the proportion of *Christensenellacea*. Their increased abundance may have reduced the degradation of the mucin layer, which in turn would decrease inflammation attributed to the translocation of bacteria and their secretions through the gut barrier.

In a study that compared the microbial composition of Japanese people ranging from infants to the elderly (26), the relative proportions of bacteria in the genera *Fusobacterium* and *Megamonas* increased with age. The positive correlation of *Fusobacterium* with fecal levels of mucin amino acids supports a previous report that showed the capacity of *Fusobacterium* species to degrade mucin (27). Odamaki et al. (26) showed a negative correlation between *Enterobacteriaceae* and a cluster of butyrate producing bacteria including *Faecalibacterium*. The decline in the proportions of *Enterobacteriaceae* in senior dogs after the consumption of the test food may have benefited the senior dogs as some of these bacteria are endotoxin producers, which compromise intestinal barrier integrity leading to inflammation. *Salmonella* belonging to *Enterobacteriaceae* also declined due to the test food consumption. Some species of *Salmonella* are major public health concerns causing salmonellosis. Although dogs are subclinical carriers of *Salmonella*, the intimate relationship between dogs and humans may lead to the risk of human exposure to *Salmonella*. The test food led to a significant relative reduction in *Salmonella* shedding by increasing the proportion of other bacteria that may have an anti-pathogenic effect.

Although not statistically significant, the relative proportions of the genera *Ruminococcus* and *Oscillospira* increased by 43% and 73%, respectively, after the consumption of the test food. Both genera belong to the family *Ruminococcaceae*, which increased significantly after the consumption of the test food. These bacteria are known to produce short-chain fatty acids (SCF) that are beneficial to the host mainly due to their anti-inflammatory effects (28, 29). They also serve as an energy source for enterocytes, regulate intestinal motility and ameliorating leaky gut syndrome (30). Bacteria in the genus *Ruminococcus* are fiber degraders and major producers of butyrate, which serves as an energy source for intestinal epithelial cells and has anti-inflammatory effects (31). Members of the genus also produce bacteriocins, which have anti-microbial effects against a wide variety of pathogenic bacteria (32).

*Oscillospira* are known to produce butyrate by relying on fermentation products secreted by other bacterial species (33). In humans, *Oscillospira* have been associated with leanness or lower body mass index in both infants and adults (34, 35). A meta-analysis by Kaakoush et al. (36), showed a negative association of the abundance of *Oscillospira* with pediatric inflammatory bowel disease. In ruminants, the abundance of *Oscillospira* is increased during the consumption of fresh green leaves and decreases upon consumption of grain containing diets (37). The presence of increased fruit and vegetable fiber in the test food may have encouraged the increase in the abundance of *Oscillospira* in the senior dogs. Conley et al. (2) showed the genus that declines the most in aged mice compared to young is *Oscillospira*. The decline in *Oscillospira* was accompanied by an increase in the marker of inflammation, monocyte chemoattractant protein-1 (MCP-1). A similar reduction in the abundance of *Oscillospira* was also associated with paracellular permeability and a decline in the anti-inflammatory cytokine, IL-10 as reported by Hamilton et al. (38).

Despite dietary levels of threonine being higher in the test food, fecal threonine declined when the dogs consumed the test food. This suggests that the increased fecal excretion of threonine is associated with the degradation of the mucin layer as reported by Weir et al, (39). The composition of the gut microbiota is a key factor in maintaining intestinal barrier integrity. The reduction in the proportion of beneficial bacteria may lead to constipation, mal-absorption and longer colonic transit time. This encourages increased presence of proteolytic bacteria, whose products of fermentation deteriorate the intestinal barrier.

The consumption of the test food reduced levels of the advanced glycation end product, pyrraline. AGE are a complex group of compounds derived from the non-enzymatic glycation of proteins, lipids, and nucleic acids. They can also be acquired from food; thus restriction of foods with high levels of AGE has been recommended to decrease circulating AGE in the body (40). AGE are known to accelerate the process of aging and they are linked to a number of age-related diseases such as diabetes, vascular and renal diseases mainly by inducing inflammation and oxidative stress (41, 42). Fecal microorganisms have been shown to be capable of degrading various AGE including pyrraline (43). The negative correlation of pyrraline with *Oscillospira*, *Christensenellaceae* and *Adlercreutzia* suggests the capability of these bacteria to degrade pyrraline or prevent its formation. The consumption of diets rich in AGE has been shown to shift the microbiota towards a more detrimental composition (44). This is in line with our correlation analyses that showed a positive association of pyrraline with *Salmonella* and *Fusobacterium*. After the consumption of the test food, the level of pyrraline in blood declined by almost the same amount (70%) as in feces. This implies induced microbial degradation of pyrraline or its precursors by the above microbes as a more likely mechanism than the test food influencing absorption of pyrraline from the GI tract. Interestingly, one of the AGE, N6-carboxymethyllysin (CML), increased in feces after the consumption of the test food. However, the level of CML in the blood did not change. This may suggest that fecal levels of CML may not be biologically significant.

The level of glycerol in the feces of the senior dogs was higher when they were fed the test food. Glycerol is known to increase water retention in the colon and thus it is used to treat constipation (45). Prolonged transit times are risks to develop various diseases due to the exposure to toxic products accumulating due to putrefaction (46). A shorter transit time leads to a limited accumulation of such products that may cause various diseases (46). People with functional constipation have been shown to contain bacteria with more abundant genes to degrade glycerol (47). The strong negative correlation between fecal levels of glycerol versus *Fusobacteria* and *Salmonella* may suggest the capacity of these bacteria to degrade glycerol. The reduction in the proportions of these bacteria after the consumption of the test food may have led to the increased levels of glycerol in feces. On the other hand, other taxa such as *Adlercreutzi*, *Christensenellaceae*, *Faecalibacterium prausnitzii*, *Ruminococcaceae*, *Phascolarctobacterium* and *Actinobacteria* correlated positively with glycerol. Weir et al. (39) found a positive association of a *Rumonococcus* species with fecal levels of glycerol and free fatty acids. To our knowledge, this study is the first to show the associations of the other taxa with fecal glycerol concentration. This may suggest the test food altered the microbial composition towards a population with higher lipase activity. Along with the increased level of glycerol in feces, the level of both fecal and plasma levels of omega fatty acids, DHA, EPA, DPA also increased in feces after the consumption of the test food due to added fish oil. Omega-3 fatty acids have health benefits throughout life by improving cardiovascular, immune, cognitive and other functions (48). Similarly, the increased level of linoleate while consuming the test food is due to the increased dietary concentration. Despite the similar levels of C:16:1 and C:18:0 fatty acids measured in the foods (Table 2), the levels of these fatty acids in feces increased after the consumption of the test food. This supports the possible increased microbial lipase activity attributed to the change in the microbial composition.

Uremic toxins are among the major metabolites that cause age-related complications. There was a marked improvement in markers of kidney health after the consumption of the test food. Circulating symmetric dimethylarginine (SDMA) has been shown to be a good biomarker of kidney function in dogs as it detects reduction in glomerular filtration rate (GFR) much earlier than serum creatinine (16). The reduction in circulating concentration of SDMA in the senior dogs after the consumption of the test food may thus indicate an increased GFR and improved kidney function.

Furthermore, the test food reduced several phenolic uremic toxins originating from microbial fermentation of protein (49). One of these metabolites was 4-ethylphenyl sulfate (4-EPS), which is also known to have a negative impact on brain health by causing anxiety-like symptoms (50, 51). The role of foods containing fruits and vegetables in improving kidney health has been reported (7, 52). The significant reduction in such metabolites in the senior dogs after the consumption of the test food may be due to the changes in the microbial composition. The two indolic uremic toxins, 5-hydroxyindole sulfate and 7-hydroxyindole sulfate, increased after the consumption of the test food. Indolic uremic toxins originate from colonic fermentation of the amino acid tryptophan (53). The test food was formulated to contain 53% higher tryptophan compared to the control 2 food. The presence of more substrate may have led to increased levels of the two indolic metabolites. However, the typical indolic uremic toxin, 3-indoxyl sulfate, was not affected by the consumption of the different diets.

In conclusion, old dogs fed fiber sources from vegetables and fruits containing high soluble fiber benefit by having a gut microbial composition promoting healthier metabolic profiles.

## Acknowledgments

E. E. G. and D. E. J designed and conducted research; M. I. J. and E.E.G. analyzed data; E.E.G. wrote the paper and had primary responsibility for final content. All authors read and approved the final manuscript. The work was funded by and performed at the Pet Nutrition Center, Hill’s Pet Nutrition, Topeka, Kansas.

